# Evolutionary divergence of potential drought adaptations between two subspecies of an annual plant: Do some constraints need to be broken?

**DOI:** 10.1101/2020.04.16.042572

**Authors:** Timothy E. Burnette, Vincent M. Eckhart

## Abstract

**Premise:** Whether mechanisms of drought adaptation tend to evolve together, evolve independently, and/or evolve constrained by genetic architecture is incompletely resolved, particularly for water-relations traits besides gas exchange. We addressed this question in two subspecies of *Clarkia xantiana* (Onagraceae), California winter annuals that diverged approximately 65,000 years ago and that are adapted, partly by differences in flowering time, to native ranges that differ in precipitation.

**Methods:** In these subspecies and in F_5_ recombinant inbred lines (RILs) from a cross between them we scored drought-adaptation traits related to phenology (times to seed germination and to flowering) and tissue water relations (measures of succulence; pressure-volume curve parameters), in common environments.

**Results:** Subspecies differed distinctly. The one native to more arid environments had some trait values associated with drought adaptation (e.g., early flowering and high succulence) but had higher osmotic potential at full turgor and lost turgor at higher water potential, indicating poorer tolerance of dehydration. Traits that differed between subspecies exhibited substantial genetic variation, with broad-sense heritability from 0.09 (stem succulence) to 0.43 (time to flowering). The genetic correlation structure suggests facilitated evolution of some trait combinations that might enhance drought adaptation (e.g., high succulence plus low turgor loss point), but the subspecies exhibit some trait combinations that do not follow genetic correlations.

**Conclusions:** As lineages diverged in their potential to escape drought by early flowering, other traits diverged as well. Genetic architecture might facilitate some correlated evolutionary responses to drought, but particular trait combinations also can evolve despite apparent genetic constraints.

---

Plant adaptation to the soil water deficits and dehydration risks associated with drought can involve diverse processes (Passioura, 1996; Tardieu, 2012; Gilbert and Medina, 2016; Volaire, 2018). For example, plants may tolerate drought by regulating stomatal conductance (Martin-StPaul et al., 2017), resisting turgor loss (Maréchaux et al., 2015), and/or having xylem that resists or recovers from hydraulic cavitation (Lens et al., 2016; Zhang et al., 2016). Where precipitation is ephemeral (e.g., some deserts) or predictably seasonal and followed by drought (e.g., Mediterranean climates), at least two other strategies can provide drought adaptation (see also (Kooyers, 2015). Plants can escape terminal drought by maturing early (Volis et al., 2002; Sherrard and Maherali, 2006), possibly assisted by rapid germination (Duncan et al., 2019a). Plants may avoid encountering drought via storing water (Ogburn and Edwards, 2012) or having seeds that do not germinate without prolonged exposure to wet conditions (Kos and Poschlod, 2010; Zeng et al., 2010; Duncan et al., 2019a; Duncan et al., 2019b). Seed dormancy can create persistent seed banks that spread the risk of encountering severe drought (and reproductive failure) among years that vary in precipitation (Cohen, 1966; Venable and Lawlor, 1980; Adondakis and Venable, 2004; Donohue et al., 2010; Saatkamp et al., 2011; Gremer and Venable, 2014). Seeking to understand how adaptation to drought evolves by these and other mechanisms is timely, given the increasing frequency and severity of climatic drought worldwide (Sheffield and Wood, 2008; Taylor et al., 2012; Keellings and Engström, 2019; Koutroulis et al., 2019).

Adaptation to drought by escape—early flowering—appears common in annuals and herbaceous perennials. Considerable evidence suggests that when and where drought arrives early, earlier flowering evolves. For example, earlier flowering is associated with xeric distributions among populations or species in *Arabidopsis* (Brassicaceae) (e.g., McKay et al., 2003; Paccard et al., 2014; Monroe et al., 2018), *Clarkia* (Vasek, 1968; Runions and Geber, 2000; Mazer et al., 2004), *Erodium* (*Geraniaceae*) (Latimer et al., 2019), *Lupinus* (Fabaceae) (Berger and Ludwig, 2014), *Mimulus* (*Erythranthe*) (Phyrmaceae) (e.g., Hall and Willis, 2006; Wu et al., 2010; Ferris and Willis, 2018; Mantel and Sweigart, 2019), *Oryza* (Poaceae) (Groen et al., 2020), and *Streptanthus* s.l. (Brassicaceae) (Pearse et al., 2019). In populations of *Brassica rapa* (Brassicaceae) (Franks, 2011) and *Mimulus laciniatus* (Dickman et al., 2019), multi-year droughts led to the evolution of earlier flowering.

When early flowering enables drought escape, how do other mechanisms of drought adaptation evolve? One pattern is a trade-off between drought escape achieved by early maturity versus drought tolerance achieved by stomatal regulation of gas exchange (Geber and Dawson, 1990; Heschel and Riginos, 2005; Emms et al., 2018). This trade-off suggests that not only might selection be relaxed for other drought adaptations if early maturity enables drought escape, but also that to achieve early maturity requires sacrificing other mechanisms of drought adaptation.

Less is known about how drought-avoiding seed-germination behavior evolves with drought escape and tolerance. The relationships may be complex, as evidenced by the fact that germination timing can alter the selective environment experienced later (Donohue, 2002). Kimball et al. (2011) found that a group of co-occurring desert winter annuals show associations between long-term demography, germination timing, flowering time, and water-use efficiency estimated by carbon-isotope discrimination. Species with consistent among-year demography tend to have early germination, early maturity (during cool weather), and high water-use efficiency, while those whose demographic performance varies widely among years tend to show contrasting traits, though with a compressed flowering period that helps escape summer drought. Dickman et al. (2019) reported more rapid seed germination, as well as earlier flowering, in

*Mimulus laciniatus* populations before and after California’s “megadrought” in the mid-2010s. As studies of tissue water relations traits—such as succulence, turgor loss point, and cavitation resistance—are rare in annuals and other herbaceous plants (Doria et al., 2019), it is even less clear how these features contribute to adaptation and/or evolve with drought escape in annuals.

How traits that confer drought escape, tolerance, and avoidance evolve in the short term will depend on the underlying genetic structure of the traits involved (Juenger, 2013). In annual plants to date there has been extensive research on genetic variation and possible constraints on models such as *Arabidopsis thaliana* (e.g., McKay et al., 2003; McKay et al., 2008; Ågren et al., 2013; Lovell et al., 2013; Mojica et al., 2016; Taylor et al., 2017; Auge et al., 2019) but much less work on other wild systems. Investigations of genetic variation and covariation in drought-related traits besides phenology and gas exchange appear to be very rare in wild annuals.

We address these knowledge gaps by investigating recent evolutionary divergence in water-relations in the California annual plant *Clarkia xantiana*. Two recognized subspecies of *C. xantiana* are lineages separated a few tens of thousands of years (Pettengill and Moeller, 2012b). They are differently adapted to geographic regions with contrasting winter and spring precipitation, partly because the subspecies from more arid areas flowers earlier (Geber and Eckhart, 2005; Benning et al., 2019). In a common environment we scored phenological and physiological traits expected to affect adaptation to drought of populations of each subspecies and of F_5_ recombinant inbred lines (RILs) from a cross between them. We found: (1) that the subspecies differ in germination phenology and tissue water relations, as well in flowering phenology; (2) that there is substantial genetic variance in these traits, plus genetic correlations that indicate major axes of trait variation; and (3) that some subspecies differences appear to have evolved in directions opposite to genetic correlations. We suggest that evolutionary change in maturation time, relative to other developmental events, may be important in understanding correlated evolution of drought adaptations in annuals.

## Materials And Methods

### Study system

*Clarkia xantiana* A. Gray occurs in mountainous areas of inland southern and central California, usually between and 500 and 1600 m elevation, on steep, sandy slopes in grassland, pine-oak savanna, and openings in chaparral (Lewis and Lewis, 1955; Eckhart and Geber, 1999). Its two recognized subspecies, *C. xantiana* ssp. *xantiana* (hereafter “*xantiana*”) and *C. xantiana* ssp. *parviflora* (Eastw.) F.H. Lewis & P.H. Raven (“*parviflora*”) separated approximately 65,000 years ago (Pettengill and Moeller, 2012b). They have mostly allopatric distributions, with *parviflora* occupying more arid areas leeward of mountain ranges that cast rain shadows (Eckhart and Geber, 1999). A narrow sympatric zone, in the Kern River drainage of the southern Sierra Nevada, represents secondary contact (Pettengill and Moeller, 2012a).

*Parviflora’s* early maturity contributes to its ability to occupy more arid regions than *xantiana*. In common environments in nature and in cultivation, *parviflora* begins to flower 1-4 weeks earlier than *xantiana* (Moore and Lewis, 1965; Eckhart and Geber, 1999; Runions and Geber, 2000; Eckhart et al., 2004). In reciprocal transplant experiments each subspecies has higher lifetime fitness (also higher than that of the other subspecies) in its native geographic range; later flowering by *xantiana* in *parviflora’* s range leads to higher mortality before setting seed, partly because of dehydration (Geber and Eckhart, 2005). In *parviflora’s* exclusive range, early flowering also increases survival by reducing the risk of being eaten by small mammals (Geber and Eckhart, 2005; Benning et al., 2019). In *xantiana*, individual performance, population growth, and geographic distribution are all strongly water-limited (Geber and Eckhart, 2005;

Eckhart et al., 2010; Eckhart et al., 2011; Kramer et al., 2011; Eckhart et al., 2017). The F_5_ RILs studied here were produced by single-seed descent from a cross between a *xantiana* population near the upper end of the lower Kern River Canyon, below Isabella Dam, and a *parviflora* population from the Pacific Crest Trail near Walker Pass, 45 km east. These are sites “8” and “47,” respectively, from previous publications (e.g., Pettengill and Moeller, 2012a). Because phenotypic variation within each RIL is almost exclusively environmental and variation among them almost exclusively genetic, among-RIL variance estimates broad-sense genetic variance in the cross, and correlations among RIL means estimate broad-sense genetic correlations underlying traits that have diverged between populations (e.g., Khasanova et al., 2019). The number of RILs was modest (14), but this feature made it feasible to estimate tissue water relations traits on several replicate individuals per RIL, from time- and labor-intensive pressure-volume curves.

### Experimental design and data collection

We cultivated parent populations and RILs in two temporal blocks, each lasting approximately 10 weeks. Seeds from the RILs came from F5 parents cultivated in a greenhouse in 2017, stored dry at room temperature for one month and at 4°C thereafter. The seeds from parental populations came from bulk collections of approximately 50 maternal families collected in natural populations in June 2018, with seeds stored dry at room temperature for one month and then dry at 4°C. With these materials we scored a series of traits based on their demonstrated and hypothesized contributions to drought adaptation (see above; Table 1).

**Table 1.**
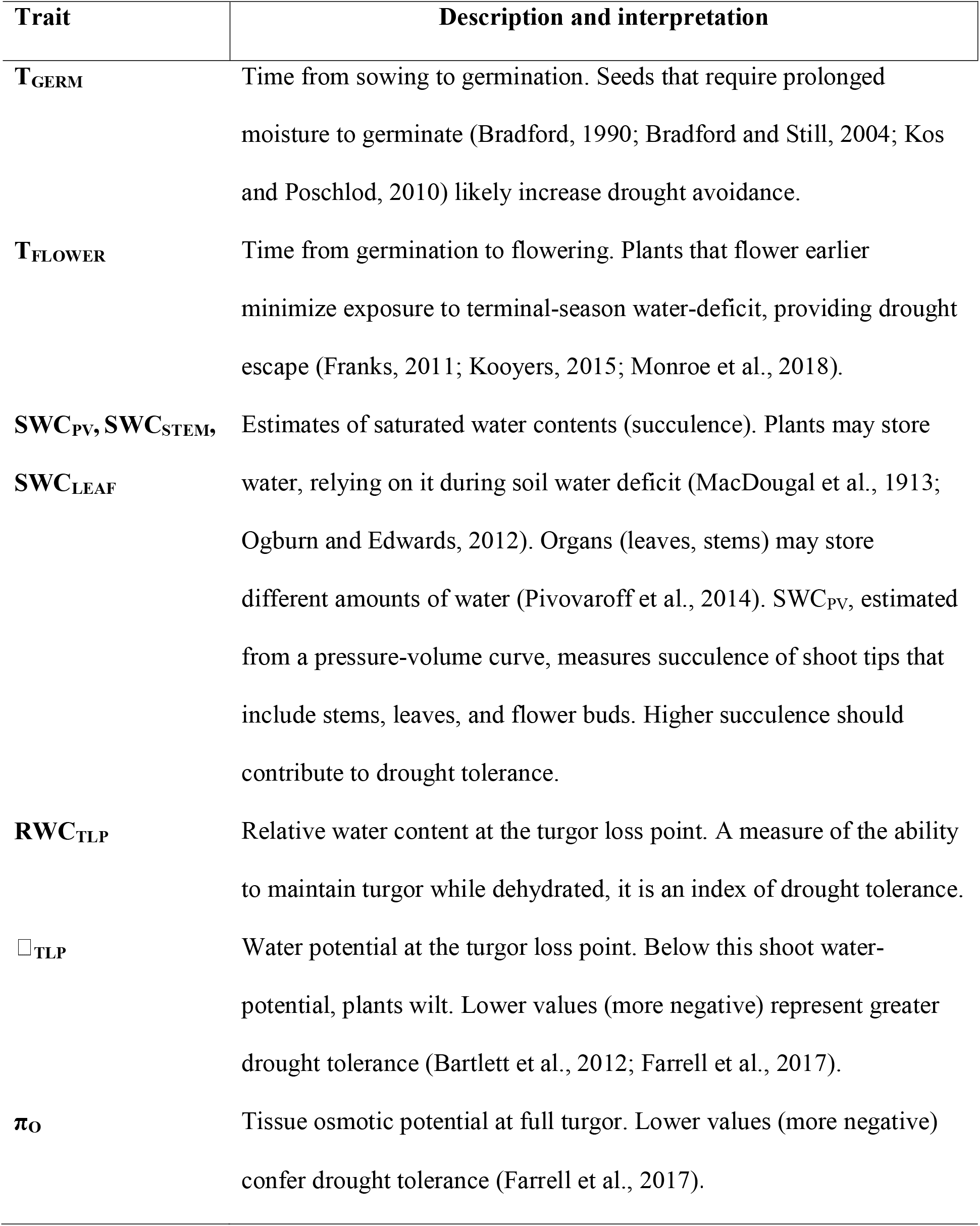
Measured traits and their interpretations in terms of drought adaptation.

Each block of the experiment began when we sowed 10 seeds per RIL and per subspecies, placing seeds into individual positions on indented seed-germination blotting paper (Seedburo, Des Plaines, Illinois, USA) inside 15 cm Petri dishes. We distributed seeds of different origin systematically (“checkerboard fashion”) into grids, to avoid genotype-environment correlation. Cotton balls placed underneath the paper, wetted with 10 ml de-ionized water, kept the paper consistently moist, and sealing with Parafilm (Bemis, Inc., Neenah, Wisconsin, USA) reduced moisture loss. After sowing we moved dishes into a growth chamber (Percival Scientific, Perry, Iowa, USA) programmed for 14 hours of dark (at 5°C) and 10 hours of light (at 15°C), simulating soil-surface conditions during natural seed germination (V. M. E., Grinnell College, unpubl.). Sampling daily, we scored each seed’s time to seed germination in days from sowing (TGERM) as radicles emerged. Terminating the seed-germination period at 14 days in each block, we found that germination was virtually complete. Only 19 of 320 seeds (6%) failed to germinate in 14 d, and we confirmed with a squeeze test (Baskin and Baskin, 2014) that 15 of these 19 were viable.

Upon its germination, we transplanted each seedling into a 13 cm wide by 8 cm deep round plastic pot filled with equal parts (v:v) potting soil (FLX, Hummert, Earth City, Missouri, USA) and fritted clay (Profile Products, Buffalo Grove, Illinois, USA). Each pot received a label with a unique code, so that in all subsequent data collection we were blind to the plant’s identity until termination of its block of the experiment. Pots resided haphazardly on two benches of a single greenhouse room. Metal-halide, high-intensity discharge lamps supplemented light for 14 hours per day, during which set points were 21°C to heat and 30°C to cool. While lights were off, set points became 13°C to heat and 18°C to cool. We applied 100 ml of fertilizer (Jack’s Classic 20-20-20 with Micronutrients, Allentown, PA) twice before flowering, and irrigation kept plants well-watered throughout. Mortality led to per-block sample sizes ranging from 2-9 individuals per RIL or parent subspecies that survived to flowering to be scored for adult traits.

After recording an individual’s time from germination to flowering (T_FLOWER_), we processed the plant for tissue water relations at the flowering stage, scoring five to six plants per day. To hydrate plants, we irrigated to saturation 1-4 hours prior to collecting data. On the top 5 cm of the central shoot (Vilagrosa et al., 2014), we measured water potentials and pressure-volume curves (Turner, 1988; Sack et al., 2011) by the “bench method,” using a pressure chamber (Model 1000, PMS Instrument Company, Albany, Oregon, USA). To accelerate desiccation after shoot tips lost turgor, we placed a fan on low speed 10 cm away from the excised shoots. We analyzed data only from those pressure-volume curves that met the following conditions: (1) initial water potential exceeded −1.00 MPa; (2) we were able to estimate water potential at seven or more points during dehydration; and (3) the curves possessed a single, obvious inflection point, indicating turgor loss. A few other anomalies (e.g., curves for which shoot water potential increased during dry-down, presumably because of measurement error) compelled us to eliminate two more curves. In all, we scored 151 plants’ pressure-volume curves, retaining 136 of them for data analysis. We used the spreadsheet tool of Sack et al. (2011) to calculate five tissue water relations parameters: (1) saturated water content at full turgor (SWC_PV_, “PV” referring to “pressure-volume” for this measure of succulence, which applies to shoot tips); (3) water potential at the turgor loss point (Ψ_TLP_); (4) osmotic potential at full turgor (π_o_); and (5) relative water content at the turgor loss point (RWC_TLP_).

For each plant, on the same day we obtained its pressure-volume curve, we estimated the mass-based succulence of its stems and leaves (Ogburn and Edwards, 2012). To estimate stem succulence (SWC_STEM_), we cut a 2 cm length of stem from each individual immediately below the shoot tip used for pressure-volume analysis, weighing it immediately and then again after 48 hours of convection drying at 65 °C. To estimate an individual’s leaf succulence (SWC_LEAF_), we recorded fresh and dry masses similarly, combining all attached, non-senescent leaves except those on the shoot tips used for pressure-volume analysis.

### Statistical analyses

We completed statistical analyses using R version 3.6.1 (R Development Core Team, 2019). We assessed normality for each variable with a Shapiro-Wilk Normality Test. For normally distributed variables we compared parent subspecies differences through a two-way type-II Wald chi-square test with subspecies (fixed) and block (random) effects using lme4 (Bates et al., 2015) and car packages (Fox and Weisberg, 2019). We used a restricted maximum-likelihood (REML) approach and derivative-free bound-constrained optimization with bobyqa. We used the lmerTest package (Kuznetsova et al., 2017) to examine block terms. Only for T_FLOWER_ was there a clear block effect (P = 0.02767, LRT = 4.8488); we ignored this, as it had negligible effects on conclusions. For days to germination, which was not normally distributed, we compared subspecies with the Scheirer-Ray-Hare extension of the Kruskal-Wallis test in the rcompanion package (Mangiafico, 2019).

For the data on RILs we estimated variance components using the lme4 (Bates et al., 2015) and insight packages (Lüdecke et al., 2019), where both RIL identity and block were treated as random. For SWC_STEM_ and Ψ_TLP_ block was singular, and therefore we ignored it. For each trait, the proportion of variance attributed to among RIL differences we interpreted as broad-sense heritability. We excluded RWC_TLP_ because it did not differ between subspecies.

To characterize the genetic correlation structure among traits, we used two approaches. First, we calculated pairwise correlations among RIL means using the corrplot package (Wei and Simko, 2017), interpreting these as broad-sense genetic correlations. Second, we looked for larger structural patterns with a principal component analysis (PCA) of RIL means, using the factoextra (Kassambara and Mundt, 2019) and Hmisc packages (Harrell and Dupont, 2019). We used singular value decomposition (SVD) to perform the PCA, with the prcomp() function. We included six traits in the analysis: T_GERM_, T_FLOWER_, Ψ_TLP_, π_0_, SWC_STEM_, and SWC_LEAF_, excluding SWC_PV_ because it correlated strongly with SWC_STEM_ and RWC_TLP_ because it did not differ between subspecies. To visualize how parent subspecies compare to RILs, we placed parents in PC1-PC2 space by multiplying z-scores for each trait and subspecies by principal component loadings.

## Results

Parent populations of the two subspecies differed distinctly in almost every trait measured (Fig. 1). Phenological traits differed in opposite directions. The evidence that seed germination differed was less clear than for other traits (H_1, 15_ = 3.10; P = 0.078,), but the trend was that *xantiana* seeds germinated in two-thirds the time of *parviflora* seeds, approximately 2 days earlier (Fig. 1a). After germination, *parviflora* flowered on average at 40 days, sooner by a third (13 days) than *xantiana* (Fig. 1b; χ^2^_1_ = 45.18; P = 1.8 x 10^-11^). In all measures of succulence *parviflora* exceeded *xantiana*, by 25% in shoot tip succulence (SWC_PV_) (Fig. 1c; χ^2^_1_ = 7.14; P = 0.0075), 25% in leaf succulence (Fig. 1d; χ^2^_1_ = 6.44; P = 0.0112), and 50% in stem succulence (Fig. 1e; χ^2^_1_ = 14.85; P = 0.00012). Water-potential traits also differed, with *xantiana* exhibiting lower averages by 0.2 MPa in π_o_ (Fig. 1g; χ^2^_1_ = 5.38; P = 0.0203) and 0.3 MPa in Ψ_TLP_ (Fig. 1h; χ^2^_1_ = 5.09; P = 0.024). RWC_TLP_ was the only variable with no evidence of a subspecies difference (Fig.1f; χ^2^_1_ = 0.53; P = 0.47), both subspecies averaging close to 80%.

**Figure 1.**
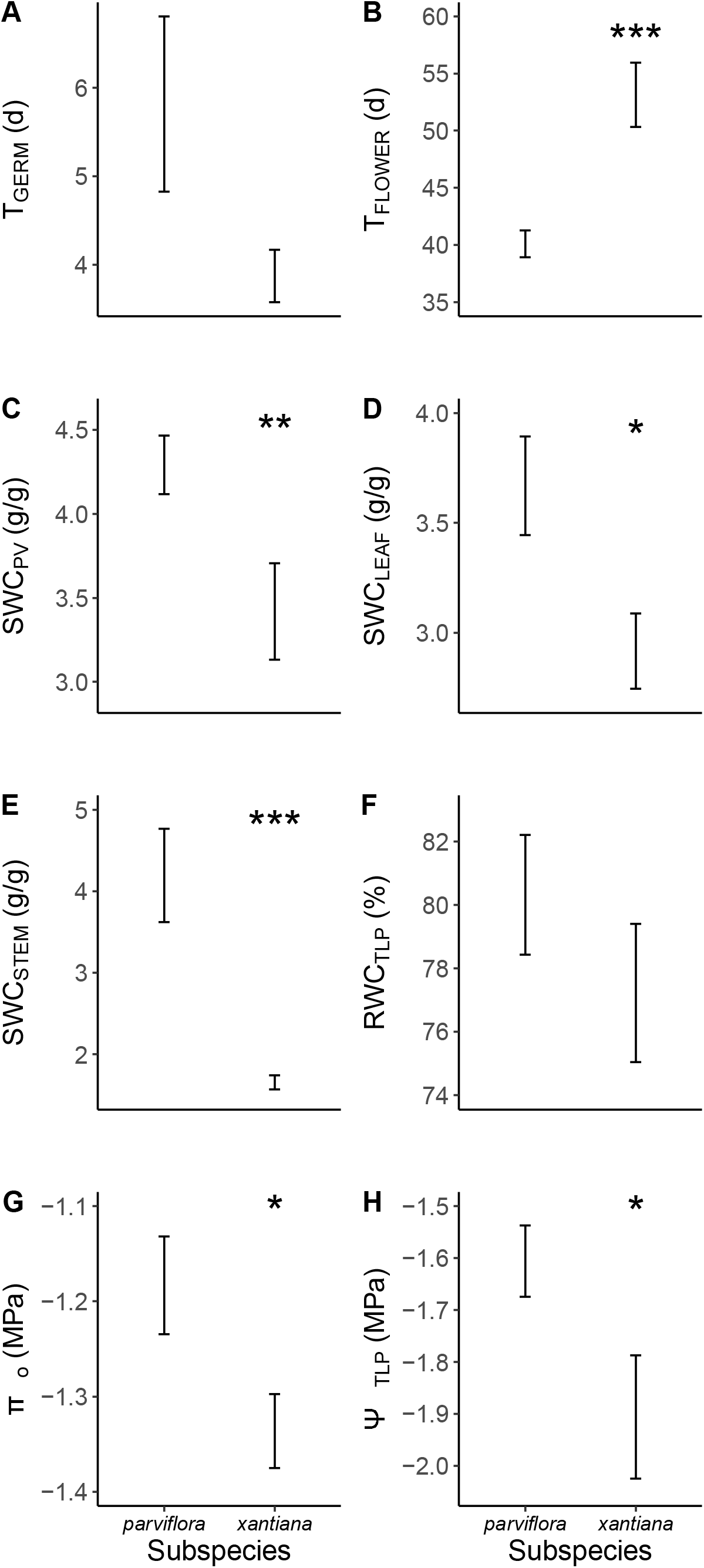
Subspecies (i.e., parent-population) comparisons for phenological and physiological traits. Symbols are means ± 1 SE. *P < 0.05, **P < 0.01, ***P < 0.001.

For every trait, the range among RILs exceeded that of the parental populations, and the among-RIL component of variance was conventionally statistically significant, indicating the presence of substantial genetic variance (Table 2; Appendix S1; see Supplemental Data with this article). Broad-sense heritability ranged from 0.09 (SWC_STEM_) to 0.42 (T_FLOWER_). Succulence measures varied approximately four-fold in heritability, suggesting somewhat independent inheritance.

**Table 2.**
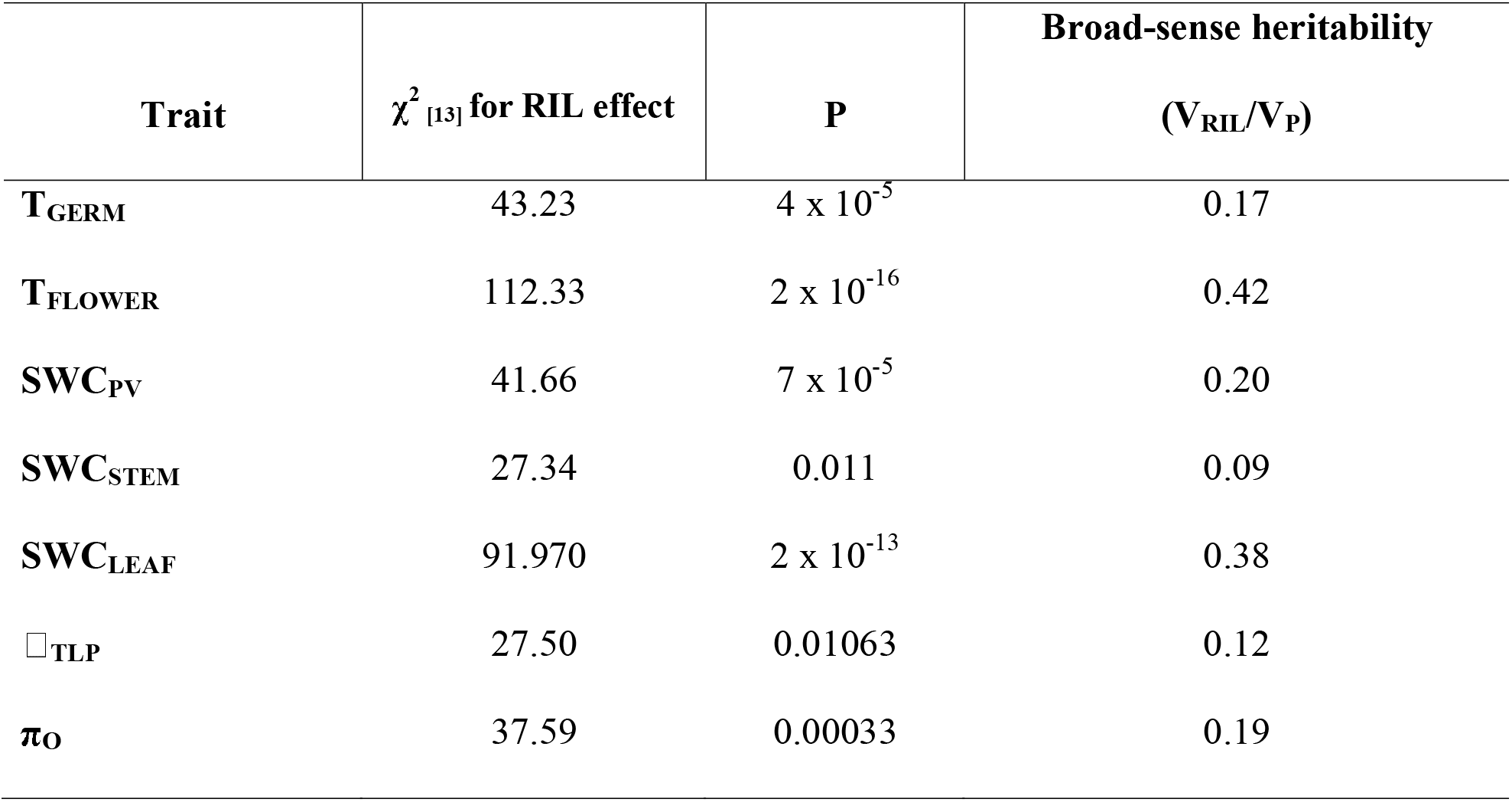
Analysis of genetic variance among 14 recombinant inbred lines for 7 water relations traits.

Six of 21 pairwise genetic correlations were strong enough to be conventionally statistically significant (Fig. 2). Strong negative correlations included those between T_FLOWER_ and both SWC_LEAF_ and SWC_PV_, between Ψ_TLP_ and both SWC_STEM_ and SWC_PV_, and between T_GERM_ and π_0_ (Fig. 2). Succulence variables correlated positively with each other, significantly so for SWC_STEM_ and SWC_PV_. Surprisingly, the two water potential variables Ψ_TLP_ and π_o_) correlated weakly (*r* = 0.19, P = 0.518).

**Figure 2.**
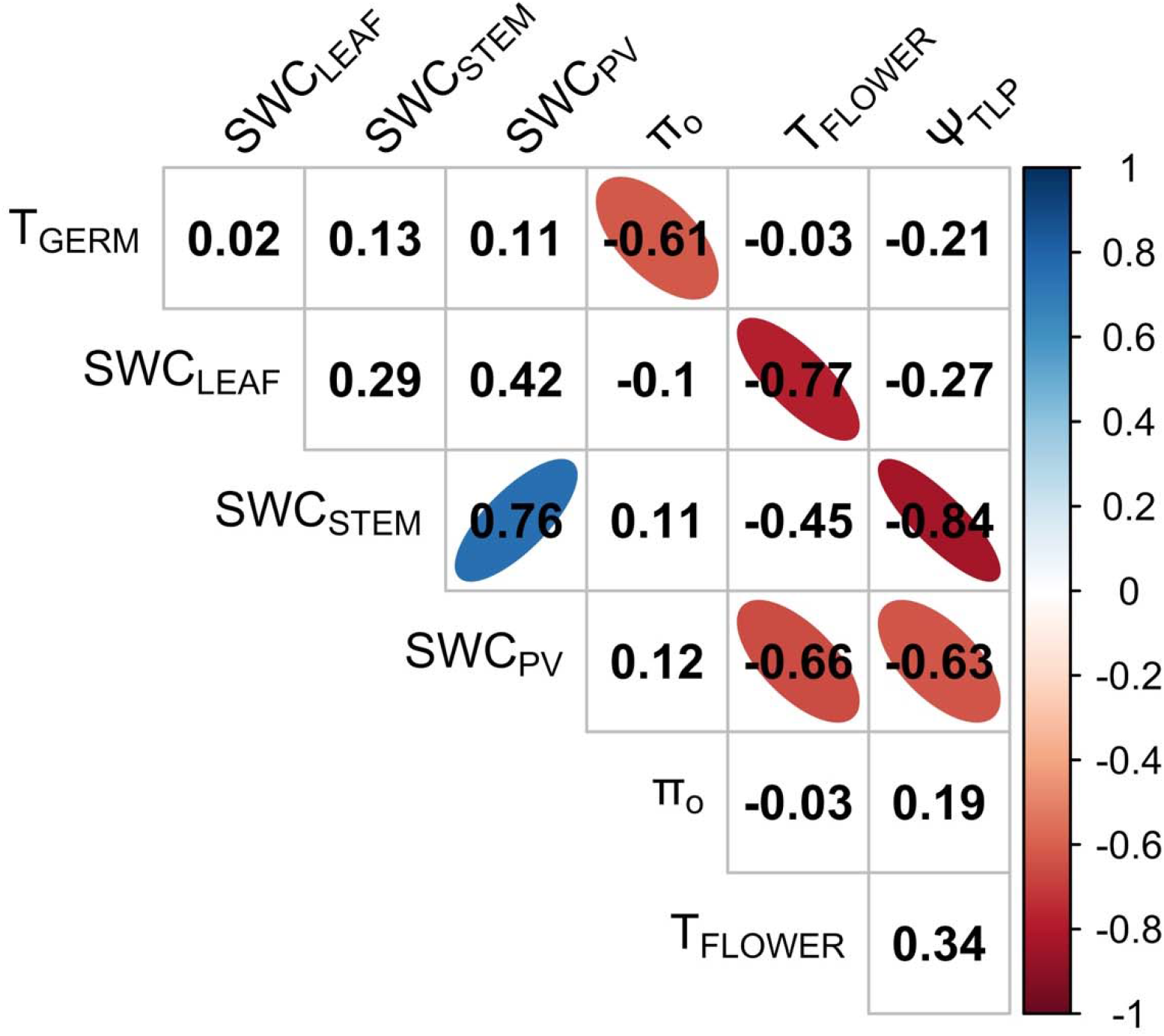
Matrix of pairwise correlation coefficients (Pearson’s *r*) among RIL means. Coefficients for which P < 0.05 (i.e., for df = 12, |*r*| > 0.532) include color- and shape-coded ellipses.

PCA of RIL means revealed two components that explained approximately 69% of the variation (Table 3; Fig. 3). The first component (PC1, accounting for 42.1%) mainly indexes T_FLOWER_ from late to early, stem and leaf succulence from low to high, and Ψ_TLP_ from high to low (Table 3; Fig. 3). The second component (PC2, accounting for 27%) mainly indexes T_GERM_ from slow to fast and π_o_ from low to high (Table 3; Fig. 3). In terms of drought adaptation mechanisms, high values of PC1 reflect drought escape (early flowering) plus components of drought tolerance (high succulence and low turgor loss point). Low values of PC2 represent genotypes that combine slow germination (drought avoidance) and low osmotic potential (drought tolerance). Plotting parent populations in the RIL-mean PC1-PC2 space shows *parviflora* as having a high value along the PC1 axis and near zero on PC2, reflecting high succulence, early flowering (and, to some extent, slow germination), despite *parviflora’s* having higher Ψ_TLP_ than *xantiana*. Meanwhile, *xantiana* positions near zero on PC1 but low on PC2, despite *xantiana* having somewhat more rapid germination than *parviflora*.

**Figure 3.**
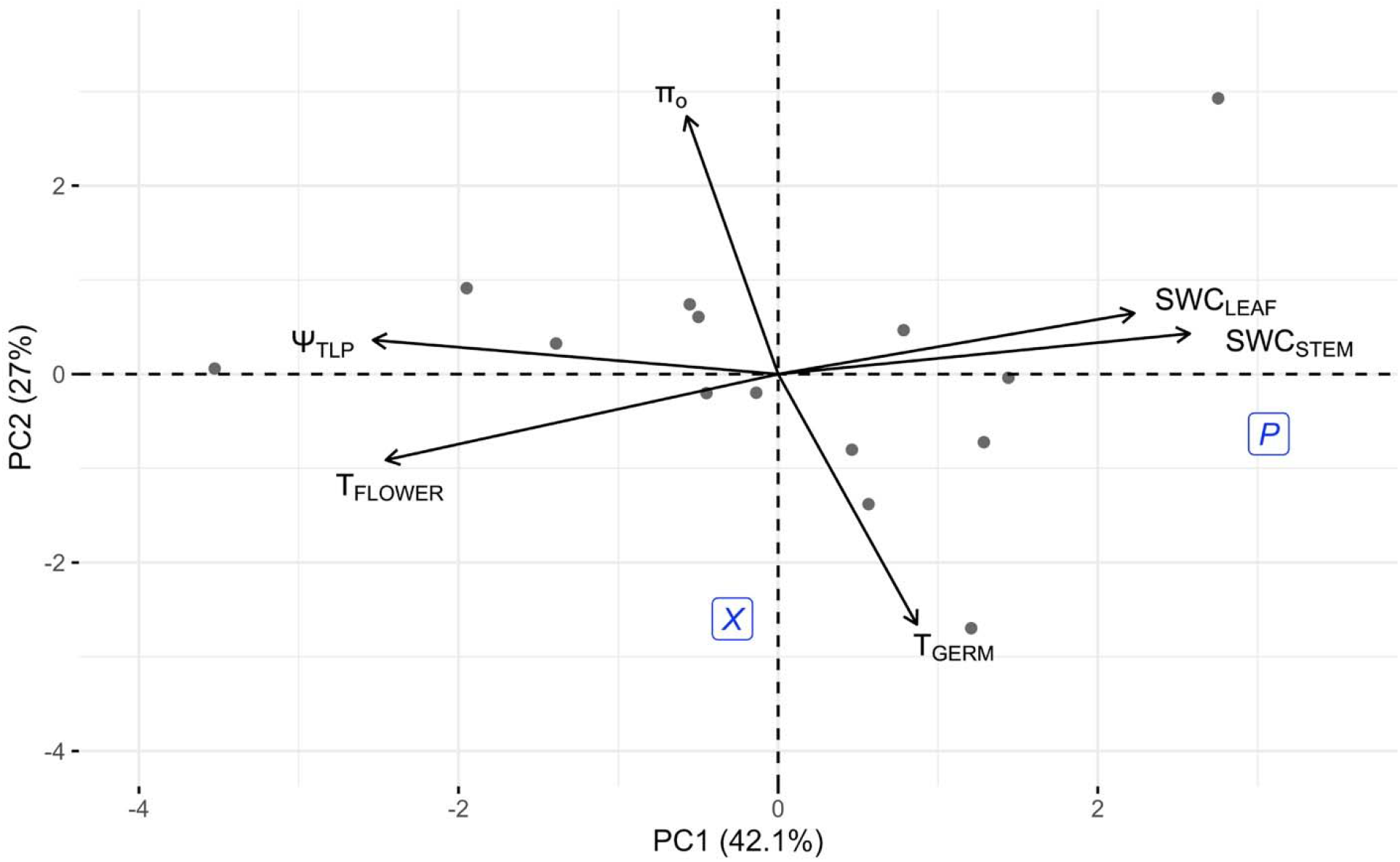
Principal components of RIL means of six phenological and physiological variables, with vectors indicating trait loadings. Points represent RILs. Subspecies’ positions, plotted in this PC1-PC2 space, appear as “*x*” and “*p*” inside boxes.

**Table 3.** Trait loadings for the principal component analysis of 14 recombinant inbred lines.

| Trait | Loading on PC1 | Loading on PC2 |
| --- | --- | --- |
| $T_{\text{GERM}}$ | 0.173 | -0.662 |
| $T_{\text{FLOWER}}$ | -0.489 | -0.227 |
| $SWC_{\text{STEM}}$ | 0.514 | 0.107 |
| $SWC_{\text{LEAF}}$ | 0.445 | 0.161 |
| $\square_{\text{TLP}}$ | -0.506 | 0.090 |
| $\pi_o$ | -0.114 | 0.682 |

## DISCUSSION

Evolutionary plant physiology provides an integrative lens to examine evolutionary tradeoffs and potential plant responses to climate change. By focusing this lens on evolutionary divergence of possible drought adaptations in *Clarkia xantiana* subspecies, we discovered that some potential drought adaptations might tend to evolve in a correlated fashion. While genetic architecture might facilitate some evolutionary responses to drought, some trait combinations of *C. xantiana* subspecies have evolved despite apparent genetic constraints.

### Subspecies differences

The subspecies differ distinctly in phenology and physiology. Previous research (Vasek, 1968; Runions and Geber, 2000; Mazer et al., 2004) documented the subspecies difference in flowering time also reported here. There is some evidence that, like other drought escapers (Geber and Dawson, 1990; Heschel and Riginos, 2005; Emms et al., 2018), *parviflora* sacrifices long-term water-use efficiency for higher rates of gas exchange, perhaps an inevitable tradeoff (Mazer et al., 2010). Here we found that *parviflora* also has higher tissue osmotic potential (π_o_) and higher turgor loss point Ψ_TLP_) than *xantiana*, tissue water relations traits that suggest sacrificed drought tolerance. Subspecies *parviflora* does not appear, however, to sacrifice all potential drought adaptations, as it has somewhat slower seed germination than *xantiana* on a saturated substrate, and it is more succulent than *xantiana* at the flowering stage. Longer time requirements for germination may confer drought avoidance, making *parviflora* less likely to germinate in places and times where soil water is insufficient to complete a life cycle (Duncan et al., 2019b) and therefore more likely to establish persistent seed banks that spread the risk of reproductive failure due to terminal drought (Saatkamp et al., 2011). Succulence may allow *parviflora* to delay, for a time, the experience of soil water deficit.

The question of whether germination timing and succulence have these benefits in *parviflora’s* native environments—independent of the benefits of drought escape—calls for field studies, such as phenotypic selection analysis (cf. Kimball et al., 2013). It is premature to interpret subspecies differences or other trait combinations as adaptive syndromes. Below we address two different questions: (1) if there were selection on pairs or suites of drought adaptations, would evolutionary responses tend to be facilitated or constrained by genetic structure; and (2) do subspecies differences indicate evolution along genetic correlations or despite genetic correlations?

### Genetic variation and evolution

RILs of a cross between the subspecies contained substantial genetic variance and covariance. Transgressive segregation appeared: wider ranges in trait values of hybrids than parents (Rieseberg et al., 1999; Devi et al., 2009; Shekoofa et al., 2013; Polania et al., 2017). Broad-sense heritability and genetic correlations estimated from RILs do not predict the precise evolutionary potential of these traits in any particular contemporary population. They are, however, a window into the nature of genetic variation that underlies subspecies differentiation, and the genetic correlations among RILs represent mostly additive genetic covariance due to pleiotropy, rather than linkage disequilibrium (Hegmann and Possidente, 1981). We interpret this variation, cautiously, as representing potential directions of drought adaptation that might be facilitated or constrained. If selection acts on a pair of traits in the same direction as their genetic correlation, there will be both direct and correlated responses, facilitating adaptation. In contrast, evolutionary responses may be constrained when selection acts orthogonally to a genetic correlation (Antonovics, 1976; but see Conner et al., 2011). Sufficiently strong selection, which can happen with large changes in drought timing (Franks et al., 2007; Groen et al., 2020), can overcome constraints, possibly facilitated by restricted recombination that preserves favorable trait combinations (Lowry and Willis, 2010; Lowry et al., 2019).

Interpreting the genetic correlations in this way (cautiously) suggests that the evolution of some combinations of drought adaptations might be facilitated. For example, mutually reinforcing responses would be expected if there were selection for early flowering (drought escape) plus high succulence (drought tolerance)—a combination that appears in *parviflora—* or if there were selection for low π_0_ (drought tolerance) and delayed germination (drought avoidance).

By visualizing genetic covariance structure among multiple traits, the principal component analysis enables a broader, though still tentative, view. PC1 suggests the potential for facilitated evolution of drought escape by early flowering, plus drought tolerance by succulence and low turgor loss point. That full combination did not, however, evolve in *parviflora*, which has a high turgor loss point. PC2 seems to capture components of drought avoidance (delayed germination) and drought tolerance (low osmotic potential at full turgor). In this way, joint responses to selection for drought avoidance by slow germination and for drought tolerance by low osmotic potential at full turgor seem likely to be facilitated. Genetic tradeoffs among drought-adaptation mechanisms clearly occur (e.g., McKay et al., 2003; Franks and Weis, 2008). In the present study, however, though the subspecies’ means have diverged in ways that suggest trade-offs, the underlying genetic variation suggests potential facilitation as well. Again, it is premature to conclude that all subspecies’ differences in traits besides flowering time represent drought adaptations favored by selection.

### Relationships among physiological traits

The present common-environment study minimized one consequence of phenological differences that applies in nature. In seasonal environments, changes in the timing of life-cycle events also change the environmental conditions experienced during those life stages, in turn affecting the expression of those traits. For example, when it is transplanted to *parviflora’s* exclusive, more arid range, *xantiana’*s flowers take longer to develop from visible buds to anthesis than they do in their native range, apparently because plants experience more severe water stress; consequently flowering occurs even later than expected (Eckhart et al., 2004). Because phenology, measured here as times to germination and flowering, generally changes environmental experiences in this way, inevitable pleiotropic changes can appear in other traits, including drought adaptations (Monroe et al., 2018; Auge et al., 2019; Takou et al., 2019). To this discussion we suggest that the pleiotropy associated with drought escape in *C. xantiana* and other annuals may arise partly from paedomorphosis: the evolution of juvenile physiology with early sexual maturity (Box and Glover, 2010). Franks and Weis (2008) found that *Brassica rapa* escapes drought via flowering at an earlier ontogenetic stage, as opposed to escaping by faster whole-plant development. The negative genetic correlation found here between flowering time and tissue succulence, with early flowering plus high succulence combination evolving in *parviflora* and late flowering plus low succulence combination in *xantiana*, might reflect age-related decline in succulence.

Finally, we point out another succulence finding that deserves investigation. Although SWC_LEAF_ and SWC_STEM_ loaded similarly along PC1 in the RIL-mean analysis, genetic correlations between these traits were not particularly strong, and within *xantiana* succulence varied considerably among organs. It is possible that the succulence of different organs can evolve more or less independently (Feng et al., 2019) and that variation in succulence within plants represents a form of hydraulic segmentation, where organs are differentially important to the water continuum (Tyree et al., 1993; Pivovaroff et al., 2014; Johnson et al., 2016). Stem succulence may play an important role in drought tolerance (Hasanuzzaman et al., 2017). The fact that *C. xantiana* stems contain gelatinous fibers in their xylem suggests specialization for water storage for terminal drought (Carlquist, 1975, 2014), but whether and how stem succulence contributes to drought adaptation in this species is unknown.

## CONCLUSIONS

As two lineages of *C. xantiana* diverged in their potential for drought escape by early flowering, other traits diverged as well, including germination behavior and tissue water relations, physiological traits that have received less attention than gas exchange in similar studies. The genetic architecture underlying subspecies might facilitate some correlated evolutionary responses to drought (e.g., high succulence and early maturity). Subspecies’ divergence followed genetic correlations in some but not all cases. Further work should address the quantitative contributions to drought adaptation of traits besides flowering time, the mechanisms of pleiotropy of flowering-time change, and the causes and consequences of organ-specific succulence.

## Acknowledgments

The authors thank Grinnell College and the National Science Foundation (DEB 1256316 & 1754157) for funding this research. We also thank the Grinnell College Greenhouse and its staff (Ashley Millet, Harper Eastman, and Marianna Cota) for caretaking of plants. John Kelly and Jamie Walters (University of Kansas) gave valuable statistical and coding advice, respectively. The David Moeller lab (University of Minnesota) and Elizabeth Queathem (Grinnell College) improved the manuscript by reviewing early drafts.

## Author contributions

T. E. B and V. M. E. conceived the study and designed the experiment. T. E. B. set up the experiment and collected the data. T. E. B. analyzed the data, with V. M. E.’s assistance. T. E. B. completed the first draft of the manuscript, and V. M. E. and T. E. B. edited and revised subsequent versions.

## Data availability statement

Pending acceptance, the raw data will be published in the Dryad data archive.

